# Development of Wheat-*Aegilops caudata* Introgression Lines and Their Characterisation Using Genome-Specific KASP Markers

**DOI:** 10.1101/2020.03.03.975326

**Authors:** Surbhi Grewal, Manel Othmeni, Jack Walker, Stella Hubbart Edwards, Cai-yun Yang, Duncan Scholefield, Stephen Ashling, Peter Isaac, Ian P. King, Julie King

## Abstract

*Aegilops caudata* L. [syn. *Ae. markgrafii* (Greuter) Hammer], a diploid wild relative of wheat (2n = 2x = 14, CC), is an important source for new genetic variation for wheat improvement due to a variety of disease resistance factors along with tolerance for various abiotic stresses. Its practical utilisation in wheat improvement can be facilitated through the generation of genome-wide introgressions leading to a variety of different wheat–*Ae. caudata* recombinant lines. In this study, we report the generation of nine such wheat–*Ae. caudata* recombinant lines which were characterized using wheat genome-specific KASP (Kompetitive Allele Specific PCR) markers and multi-colour genomic *in situ* hybridization (mcGISH). Of these, six lines have stable homozygous introgressions from *Ae. caudata* and will be used for future trait analysis. Through a combination of molecular and cytological analysis of all the recombinant lines, we were able to physically map 182 KASP markers onto the seven *Ae. caudata* chromosomes, of which 155 were polymorphic specifically with only one wheat subgenome. Comparative analysis of the physical positions of these markers in the *Ae. caudata* and wheat genomes confirmed that the former had chromosomal rearrangements with respect to wheat, as previously reported. These wheat–*Ae. caudata* recombinant lines and KASP markers provide a useful genetic resource for wheat improvement with the latter having a wider impact as a tool for detection of introgressions from other Aegilops species into wheat.

## INTRODUCTION

Wheat is one of the most widely cultivated crops worldwide, contributing about a fifth of the total calories and protein consumed by humans. While wheat production needs to increase to feed the ever-growing population, wheat yields are plateauing due to the relatively small amount of genetic variation available in the gene pool of wheat that can be used to develop new high yielding varieties adapted to the changing environment. However, wild relatives of wheat possess genetic diversity that could be exploited in wheat breeding programmes through introgression of small segments of their genome, carrying desirable traits, into wheat (Feuillet et al., 2008).

Aegilops species are an important source of such genetic variation required for the improvement of cultivated bread wheat, *Triticum aestivum* (2n = 6x = 42, AABBDD). *Aegilops caudata* L. [syn. *Ae. markgrafii* (Greuter) Hammer] is a diploid wild relative of wheat (2n = 2x = 14, CC) native to the north-eastern Mediterranean basin with the main distribution from Greece to northern Iraq (Ohta and Yasukawa, 2015). This species is the C-genome progenitor of the allotetraploids *Aegilops cylindrica* (2n = 4x = 28; DDCC) and *Aegilops truncialis* (2n = 4x = 28; UUCC). *Ae. caudata* carries resistance genes for several diseases (Makkouk et al., 1994), especially against stripe rust (Baldauf et al., 1992; Toor et al., 2016; Valkoun et al., 1985), leaf rust (Gong et al., 2017; Iqbal et al., 2007; Riar et al., 2012), stem rust (Dyck et al., 1990; Gong et al., 2017) and powdery mildew (Baldauf et al., 1992; Gill et al., 1985; Valkoun et al., 1985; Weidner et al., 2012) and resistance against pests such as greenbug and hessian fly (Baldauf et al., 1992; Gill et al., 1985). Moreover, some accessions of *Ae. caudata* have shown freezing tolerance (Barashkova and Migushova, 1984; Iriki et al., 2001), salt tolerance (Gorham, 1990) and increased grain mineral content (Wang et al., 2011). Thus *Ae. caudata* provides an important source of genetic variation for agronomically important traits that can be transferred into wheat as has been previously done for other Aegilops species (King et al., 2018; King et al., 2017).

In the past, the lack of information available for the cytomolecular identification of the C-genome chromosomes has hampered the use of the genetic potential of *Ae. caudata* in wheat pre-breeding programmes. In addition, poor knowledge of syntenic relationships between wheat and *Ae. caudata* chromosomes has been another obstacle hampering the use of its genetic diversity in wheat breeding. Over the past decades, however, efforts have been made to study the molecular organisation of the *Ae. caudata genome* and its homology with wheat homoeologous groups. Friebe et al. (1992) showed that *Ae. caudata* has a highly asymmetric karyotype distinct from the metacentric and submetacentric chromosomes of most Triticeae species suggesting the distortion of chromosome collinearity compared to wheat. Fluorescence *in situ* hybridisation (FISH) and molecular marker analysis of flow-sorted C-genome chromosomes confirmed genome rearrangements (Molnár et al., 2016; Molnár et al., 2015). A set of wheat (cv. Alcedo)-*Ae. caudata* addition lines B-G (Schubert and Blüthner, 1992; Schubert and Blüthner, 1995) have also been characterised extensively in previous studies using cytogenetic markers (Friebe et al., 1992), isozyme analysis (Schmidt et al., 1993), Simple Sequence Repeat (SSR) markers (Gong et al., 2017; Niu et al., 2018; Peil et al., 1998), FISH with cDNA probes (Danilova et al., 2017), Conserved Orthologous Sequence (COS) and PCR-based Landmark Unique Gene (PLUG) markers (Gong et al., 2017), and sequential FISH and genomic *in situ* hybridization (GISH) (Niu et al., 2018). Many of these studies found that the Alcedo-*Ae. caudata* addition lines carried several inversions and translocations although the assignment of *Ae. caudata* chromosomes to corresponding wheat homoeologous groups was not consistent between them.

Despite numerous studies using wheat-*Ae. caudata* addition lines and the development of molecular markers capable of detecting *Ae. caudata* chromosomes in a wheat background, its utilisation for wheat improvement through recombination with wheat chromosomes has rarely been reported (Dyck et al., 1990; Iqbal et al., 2007; Riar et al., 2012; Toor et al., 2016; Weidner et al., 2012). In this work, we report the generation of six stable wheat-*Ae. caudata* recombinant lines and the physical location of 182 KASP markers on *Ae. caudata* chromosomes, that can be used to detect the presence of *Ae. caudata* introgressions in a wheat background.

## MATERIALS AND METHODS

### Plant Material

*Triticum aestivum* cv. Paragon *ph1/ph1* mutant (2n = 6x = 42) was pollinated with *Ae. caudata* accession 2090001 (obtained from Germplasm Resource Unit, GRU at the John Innes Centre, Norwich, UK) to produce F_1_ interspecific hybrids. The origin, according to the GRU database, Seedstor, of accession 2090001 is unknown. There is no trait data available for this accession and it was thus, chosen at random. *T. aestivum* cv. Alcedo-*Ae. caudata* addition lines B-G (TA3558-TA3563), and the Alcedo cultivar were obtained from Wheat Genetics Resource Center (WGRC) at Kansas State University, USA.

After being grown to maturity, the F_1_ wheat -*Ae. caudata* hybrids were used as the female parent and backcrossed with Paragon wheat, carrying the wild-type *Ph1* locus intact, to generate a BC_1_ population. The BC_1_ individuals were then recurrently pollinated with Paragon *Ph1/Ph1* to produce BC_2_, BC_3_ and BC_4_ lines. Three heads from each plant in each backcross population were bagged to allow self-fertilisation.

### Genotyping with KASP^TM^ markers

Genomic DNA was isolated from leaf tissue of 10-day old seedlings in a 96-well plate as described by Thomson and Henry (1995). All back-crossed lines were genotyped alongside five wheat genotypes (Chinese Spring, Paragon, Pavon, Highbury and Alcedo) and the *Ae. caudata* accession as controls. The wheat (Alcedo)-*Ae. caudata* B-G addition were also used as control lines to verify the specificity of the KASP markers (Grewal et al., 2020) to each of the *Ae. caudata* chromosomes.

For each KASP™ marker, two allele-specific forward primers and one common reverse primer were used (Supplemental Table 1). Genotyping reactions were performed in a ProFlex PCR system (Applied Biosystems by Life Technology) in a final volume of 5 µl with 1 ng genomic DNA, 2.5 µl KASP reaction mix (ROX), 0.068 µl primer mix and 2.43 µl nuclease free water. PCR conditions were set as 15 min at 94°C; 10 touchdown cycles of 10 s at 94°C, 1 min at 65– 57°C (dropping 0.8°C per cycle); and 35 cycles of 10 s at 94°C, 1 min at 57°C.

**Table 1.** Assignment of genome-specific and genome-nonspecific KASP markers, and consequently homoeologous groups, to *Ae. caudata* chromosomes derived from Alcedo-*Ae. caudata* disomic addition lines and the distribution of the genome-specific KASP markers on the wheat A-, B- and D-genomes.

| *Ae. caudata*<br>chromosome | Alcedo-*Ae.*<br>*caudata*<br>addition<br>line | Genome-specific KASP<br>markers |  |  | Genome-nonspecific<br>markers | Total<br>number of<br>KASP<br>markers | Homoeologous<br>group(s) |
| --- | --- | --- | --- | --- | --- | --- | --- |
|  |  | A | B | D |  |  |  |
| 1C | - | 7 | 7 | 9 | 3 | 26 | 1 |
| 2C | B | 10 | 6 | 10 | 1 | 27 | 2/4 |
| 3C | F | 2 | 6 | 7 | 7 | 22 | 3 |
| 4C | G | 4 | 9 | 6 | 4 | 23 | 4/2/3/7 |
| 5C | C | 8 | 14 | 10 | 1 | 33 | 5 |
| 6C | D | 3 | 7 | 5 | 8 | 23 | 6 |
| 7C | E | 6 | 4 | 15 | 3 | 28 | 7 |
| **Total** |  | **40** | **53** | **62** | **27** | **182** |  |

Fluorescence detection of the reactions was performed using a QuantStudio 5 (Applied Biosystems) and the data analysed using the QuantStudio™ Design and Analysis Software V1.5.0 (Applied Biosystems).

### Multi-colour Genomic *In Situ* Hybridisation (mcGISH)

Preparation of the root-tip metaphase chromosome spreads, the protocol for mcGISH and the image capture was as described in (Grewal et al., 2020). In summary, genomic DNAs from young leaves of the three putative diploid progenitors of bread wheat, i.e. *Triticum urartu* (A-genome), *Aegilops speltoides* (B-genome), and *Aegilops tauschii* (D-genome) and of *Ae. caudata*, were isolated using extraction buffer [0.1M Tris-HCl (pH 7.5), 0.05M EDTA (pH 8.0), 1.25% SDS]. Samples were incubated at 65°C for 1 h before being placed on ice and mixed with ice cold 6M NH_4_C_2_H_3_O for 15 mins. The samples were then spun down, the supernatant mixed with isopropanol to pellet the DNA and the isolated DNA further purified with phenol/chloroform. The genomic DNA of 1) *T. urartu* was labelled by nick translation with ChromaTide™ Alexa Fluor™ 488-5-dUTP (Invitrogen; C11397; coloured green), 2) *Ae. speltoides* was labelled by nick translation with DEAC-dUTP (Jena Bioscience; NU-803-DEAC; coloured blueish purple), 3) *Ae. tauschii* was labelled with ChromaTide™ Alexa Fluor™ 594-5-dUTP (Invitrogen; C11400; coloured red) and 4) *Ae. caudata* was labelled by nick translation with ChromaTide™ Alexa Fluor™ 546-14-dUTP (Invitrogen; C11401; coloured yellow). Slides were probed with the labelled genomic DNAs in the ratio 3:3:6:1 (green: blue: red: yellow). All slides were counterstained with DAPI and analysed using a high-throughput, fully automated Zeiss Axio ImagerZ2 upright epifluorescence microscope (Carl Zeiss Ltd, Oberkochen, Germany) with filters for DAPI (blue), Alexa Fluor 488 (green), Alexa Fluor 594 (red), Alexa Fluor 546 (yellow) and DEAC (aqua). Photographs were taken using a MetaSystems Coolcube 1m CCD camera. Further slide analysis was carried out using Metafer4 (automated metaphase image capture) and ISIS (image processing) software (Metasystems GmbH, Altlussheim, Germany).

### Fluorescence *In Situ* Hybridisation (FISH)

For fluorescence *in situ* hybridization (FISH), two repetitive DNA sequences pSc119.2 (McIntyre et al., 1990), and pAs.1 (Rayburn and Gill, 1986) were labelled with Alexa Fluor 488-5-dUTP (green) and Alexa Fluor 594-5-dUTP (red), respectively, and hybridized to the slide in the ratio 2:1 (green: red). Subsequent counterstaining and image capture were performed as described for GISH. The wheat FISH karyotype used for assigning chromosomes was that established by Tang et al. (2014).

## RESULTS

### Assignment of KASP^TM^ Markers to *Ae. caudata* Chromosomes

In a previous study, we reported the generation of 183 KASP markers polymorphic between hexaploid wheat and *Ae. caudata* accession 2090001 (Grewal et al., 2020). Although, the physical location of these markers on the wheat genome was reported, the study did not address the physical location of these markers on the *Ae. caudata* genome.

In this work, the KASP markers were tested on the Alcedo-*Ae. caudata* addition lines carrying chromosomes B-G and the assignment of *Ae. caudata* chromosomes to homoeologous groups was determined based on the distribution of the KASP markers among the addition lines. When a KASP marker detected the presence of *Ae. caudata* in any one of the addition lines, the marker was assigned to the corresponding C-genome chromosome. As shown in Table 1, 182 out of the 183 KASP markers were individually assigned to an *Ae. caudata* chromosome. An addition line for chromosome A is absent from this set, but reports have suggested it to be homoeologous to group 1 of wheat (Danilova et al., 2017; Friebe et al., 1992; Niu et al., 2018). Thus, the 26 markers that did not detect *Ae. caudata* in the addition lines B-G were assigned to chromosome 1C.

There were 155 KASP markers specific to a wheat subgenome and 27 which had more than one corresponding sequence in the wheat genome and hence, were genome-nonspecific for wheat (Supplementary Table 2). In total, 40 KASP markers were developed for the C-genome of *Ae. caudata* that were polymorphic with the A-genome of wheat, 53 with the B-genome and 62 with the D-genome of wheat. Chromosome 5C had the maximum number of KASP markers assigned to it (33) whereas the rest of the C-genome chromosomes had between 22 and 28 KASP markers assigned to them (Table 1). However, some C-genome chromosomes such as 2C and 4C, had markers assigned from non-homoeologous groups as shown in Table 1.

### Identification of Wheat-*Ae. caudata* Recombinant Lines

The KASP markers were also used to genotype a set of 181 wheat-*Ae. caudata* BC_x_F_y_ lines to detect the presence of *Ae. caudata* introgressions in wheat. This set included 3 BC_1_, 7 BC_2_, 21 BC_3_, 15 BC_3_F_1_, 7 BC_3_F_2_, 42 BC_4_, 47 BC_4_F_1_ and 39 BC_5_ plants. The genotyping analysis indicated nine advanced back-crossed lines that had one or two recombinant chromosomes without the presence of any whole *Ae. caudata* chromosome (Table 2). Two recombinants were obtained from chromosome 1C, one from chromosome 4C, four from chromosome 5C and one from chromosome 7C. One introgression line had two of the above-mentioned recombinant chromosomes, one from 1C and one from 5C.

**Table 2.** List of wheat-*Ae. caudata* introgression lines containing recombinant chromosomes (RC) obtained in this study, the copy number of the RC(s) and the total number of chromosomes (2n) in each line.

| Introgression<br>Line | Recombinant chromosome(s)<br>(RC) | Copies<br>of RC(s) | Total<br>chromosome<br>number 2n |
| --- | --- | --- | --- |
| BC<sub>4</sub>F<sub>1</sub>-157-9 | T1AL-1CL.1CS | 2 | 42 |
| BC<sub>4</sub>F<sub>1</sub>-157-2 | T5AL-5CL.5CS#1 | 2 | 42 |
| BC<sub>4</sub>F<sub>1</sub>-158-12 | T5AL-5CL.5CS#2 | 2 | 42 |
| BC<sub>4</sub>F<sub>1</sub>-162-6 | T5DL-5CL.5CS | 2 | 42 |
| BC<sub>3</sub>F<sub>2</sub>-160-4 | T7DL-7CL.7CS | 2 | 41 |
| BC<sub>4</sub>F<sub>1</sub>-156-3 | T1AL-1CL.1CS, T5AL-5CL.5CS#2 | 2 | 42 |
| BC<sub>4</sub>-284-4 | T1DS.1DL-1CL | 1 | 42 |
| BC<sub>4</sub>F<sub>1</sub>-164-2 | T3DS.3DL-4CL | 1 | 41 |
| BC<sub>3</sub>F<sub>1</sub>-401-8 | T5BL-5CL.5CS | 1 | 42 |

After self-fertilisation, six of these BC_x_F_y_ lines showed stable homozygous introgressions which were validated by mcGISH analysis (Figures 1a-f). Five out of these six lines had 42 chromosomes with the sixth line having 41 chromosomes including the recombinant chromosomes (Table 2). However, prior to mcGISH analysis, the genome-specific KASP markers enabled the identification of the homozygous lines and the wheat chromosome that had recombined with the *Ae. caudata* segment in the homozygous recombinant lines (Figures 2a-h).

**Figure 1.**
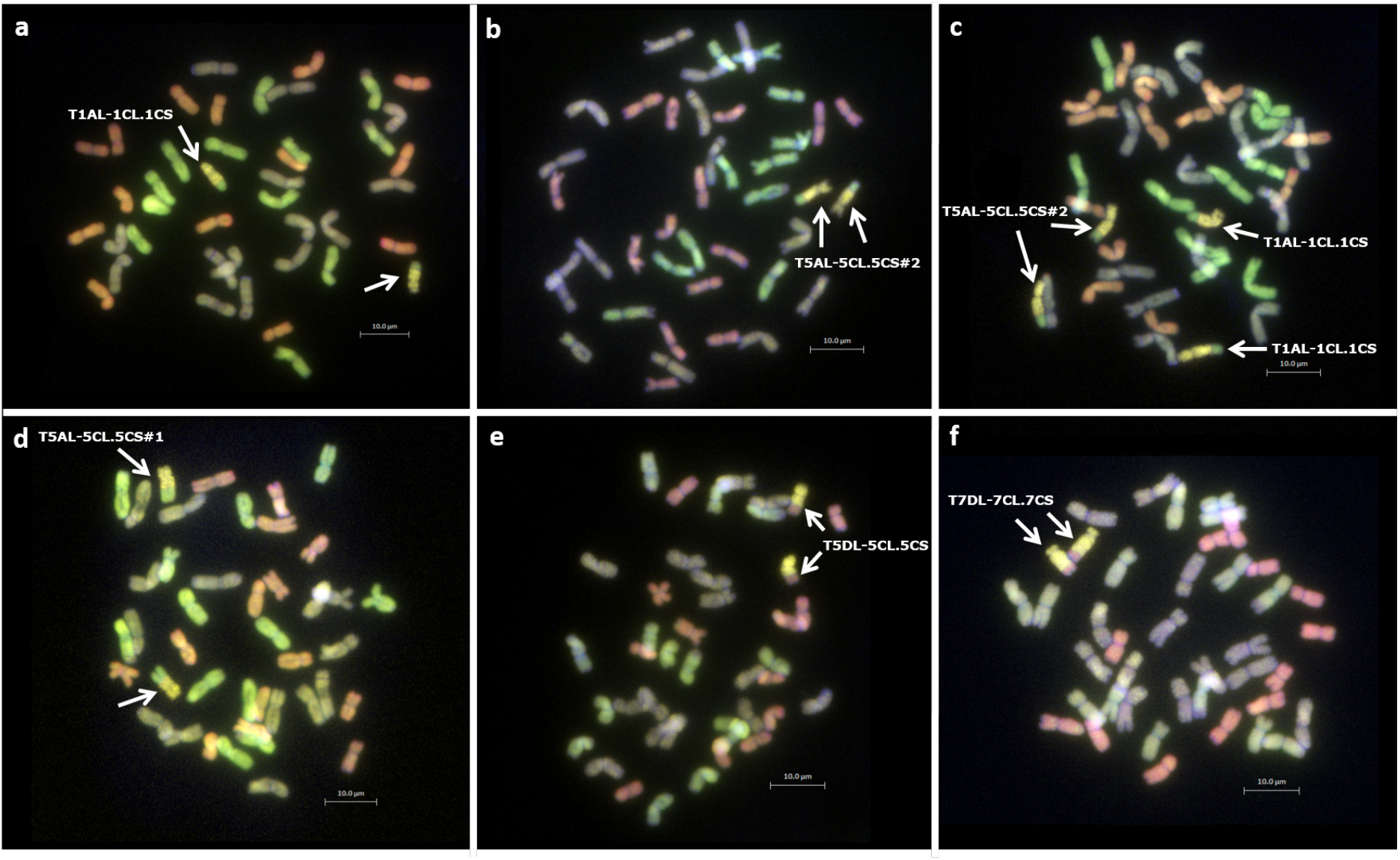
Identification of wheat-*Ae. caudata* recombinant chromosomes in homozygous lines through mcGISH analysis of root-tip metaphase chromosomes spreads. Arrows indicate recombinant chromosomes. **a)** BC_4_F_1_-157-9, **b)** BC_4_F_1_-158-12, **c)** BC_4_F_1_-156-3, **d)** BC_4_F_1_-157-2, **e)** BC_4_F_1_-162-6 and **f)** BC_3_F_2_-160-4. *Ae. caudata* segments (yellow) recombined with the A-(green) and D-(red) genome chromosomes of wheat. Wheat B-genome is represented as greyish/purple chromosomes.

**Figure 2.**
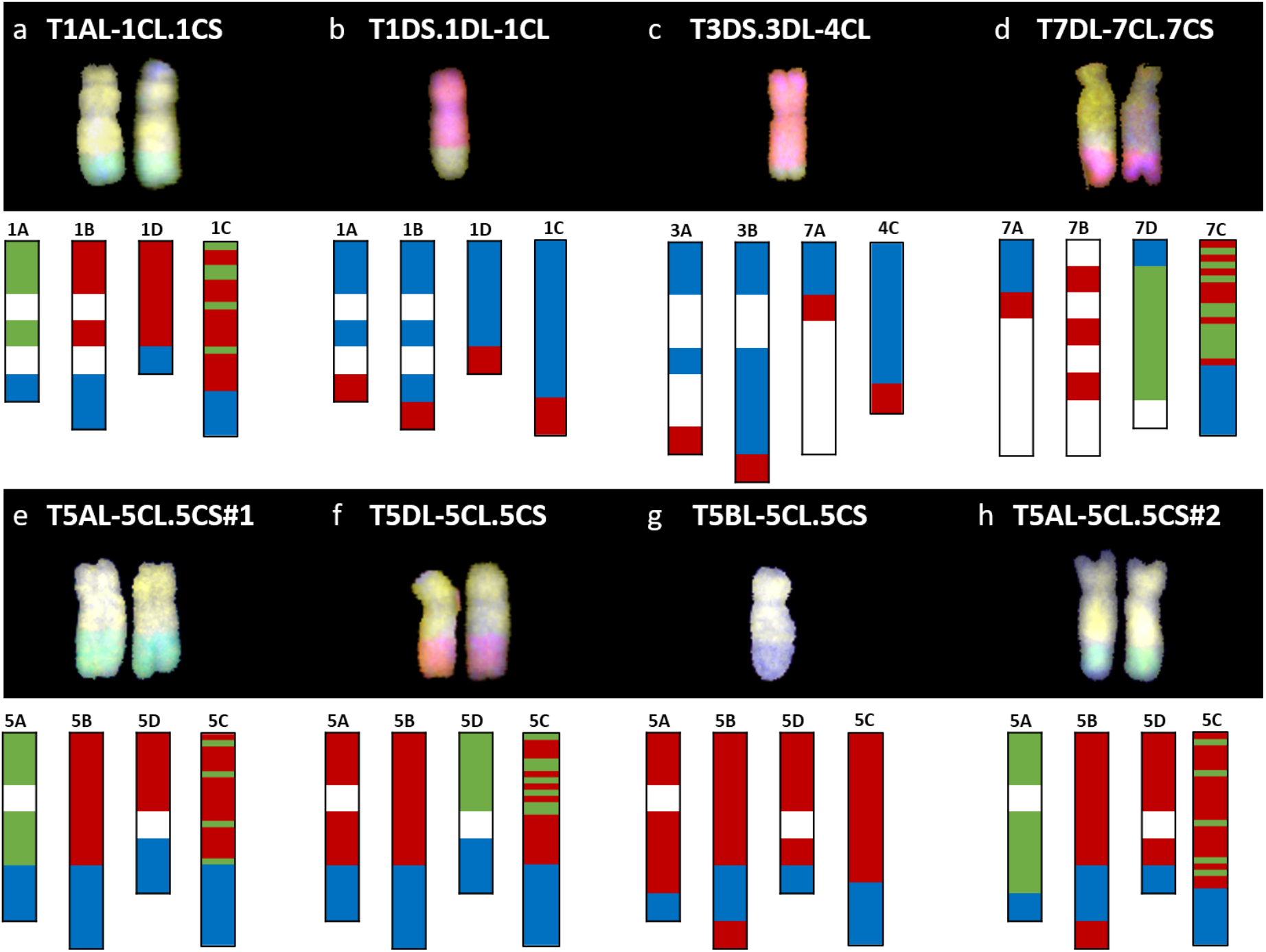
Molecular characterisation of wheat-*Ae. caudata* introgression lines using KASP markers and mcGISH. Top: the mcGISH image of the recombinant chromosome(s) and bottom: the genotyping data with KASP markers, of 8 wheat-*Ae. caudata* introgression lines. **a)** BC_4_F_1_-157-9, **b)** BC_4_-284-4, **c)** BC_4_F_1_-164-2, **d)** BC_3_F_2_-160-4, **e)** BC_4_F_1_-157-2, **f)** BC_4_F_1_-162-6, **g)** BC3F1-401-8, and **h)** BC_4_F_1_-158-12. C-genome segment is represented in yellow, A-genome in green, B-genome in blue and D-genome in red. The genotyping is displayed with the markers distributed on the wheat chromosomes and the same set of markers ordered on the *Ae. caudata* chromosomes. White areas on the wheat chromosomes indicate the absence of KASP markers. Coloured regions represent different genotype calls. Blue represents KASP markers homozygous for the wheat allele, red represents a heterozygous call and green represents a homozygous call for the *Ae. caudata* allele.

Figure 2 shows the mcGISH image of the recombinant chromosome(s) (top) and the KASP marker analysis (bottom) of 8 wheat-*Ae. caudata* introgression lines. The mcGISH is displayed with the C-genome segment in yellow, A-genome in green, B-genome in blue and D-genome in red. The genotyping is displayed with the markers physically located on the wheat chromosomes (for e.g., chromosomes 1A, 1B and 1D in Figure 2a) and the same set of markers on the *Ae. caudata* chromosomes (for e.g., chromosome 1C in Figure 2a). The white areas on the wheat chromosomes indicate the absence of KASP markers polymorphic with *Ae. caudata* whereas the coloured regions represent different genotype calls. The blue represents where the KASP markers were called homozygous for the wheat allele, the red represents a heterozygous call and the green represents a homozygous call for the *Ae. caudata* allele.

When a line had a homozygous introgression, the KASP markers on the wheat chromosome involved in the recombination showed homozygous green calls as shown in Figures 2a, 2d, 2e, 2f, and 2h. This is due to the genome-specificity of the KASP markers in wheat. Thus, when two copies of the *Ae. caudata* segment replaced both copies of the homoeologous wheat regions in the homozygous recombinant chromosomes, it resulted in no wheat allele for the genome-specific KASP markers in those regions. All other KASP markers detecting the *Ae. caudata* segment but not physically located on the recombinant chromosome show a heterozygous call to indicate the presence of *Ae. caudata* in a wheat background. The marker analysis in Figures 2a, 2e and 2h indicated the C-genome segment had recombined with the A-genome of wheat whereas in Figures 2b and 2f, the markers indicated the C-genome segment had recombined with the D-genome of wheat. The mcGISH analysis, above, confirmed these results.

When a line had a heterozygous introgression, as shown in Figures 2b, 2c and 2g, all the KASP markers that were located within the introgressed *Ae. caudata* segment gave a heterozygous call and thus, were not able to indicate which wheat chromosome the segment had recombined with. The mcGISH analysis (top), however, indicated the wheat genome, B (Figure 2g) or D (Figures 2b and 2c), present in these recombinant chromosomes.

### Physical Ordering of KASP Markers on C-Genome Chromosomes and Comparative Analysis with Wheat

Based on the molecular marker analysis of the Alcedo-*Ae. caudata* addition lines (Table 1) and the wheat-*Ae. caudata* back-crossed population (Figure 2; bottom), the mcGISH analysis of the recombinant chromosomes (Figure 2; top) and previous reports on C-genome chromosomal rearrangements (Danilova et al., 2017), the 182 KASP makers were tentatively ordered onto the seven *Ae. caudata* C-genome chromosomes (Figure 3). A comparative analysis of the markers on the C-genome chromosomes and their physical positions on the wheat A, B and D-genome chromosomes (Supplemental Table 2), as indicated by the wheat reference genome assembly RefSeqv1 (IWGSC et al., 2018), is shown in Figure 4.

**Figure 3.**
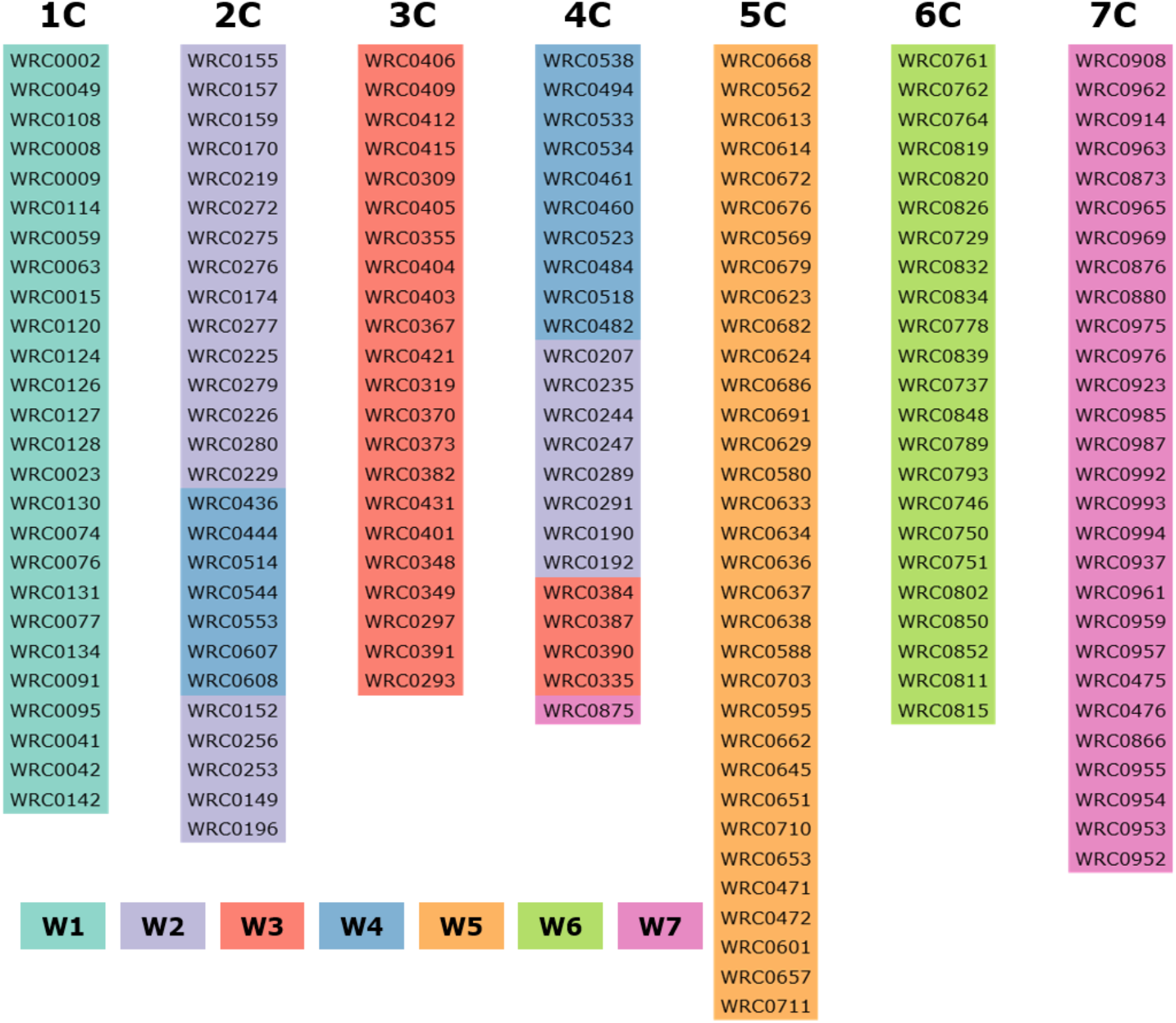
Physical order of the 182 KASP markers assigned specifically to each *Ae. caudata* chromosome. W means wheat and W1–W7 represents wheat homoeologous group 1–7, respectively. Chromosomes 2C showing markers from homoeologous group 2/4 and chromosome 4C showing markers from homoeologous groups 4/2/3/7.

**Figure 4.**
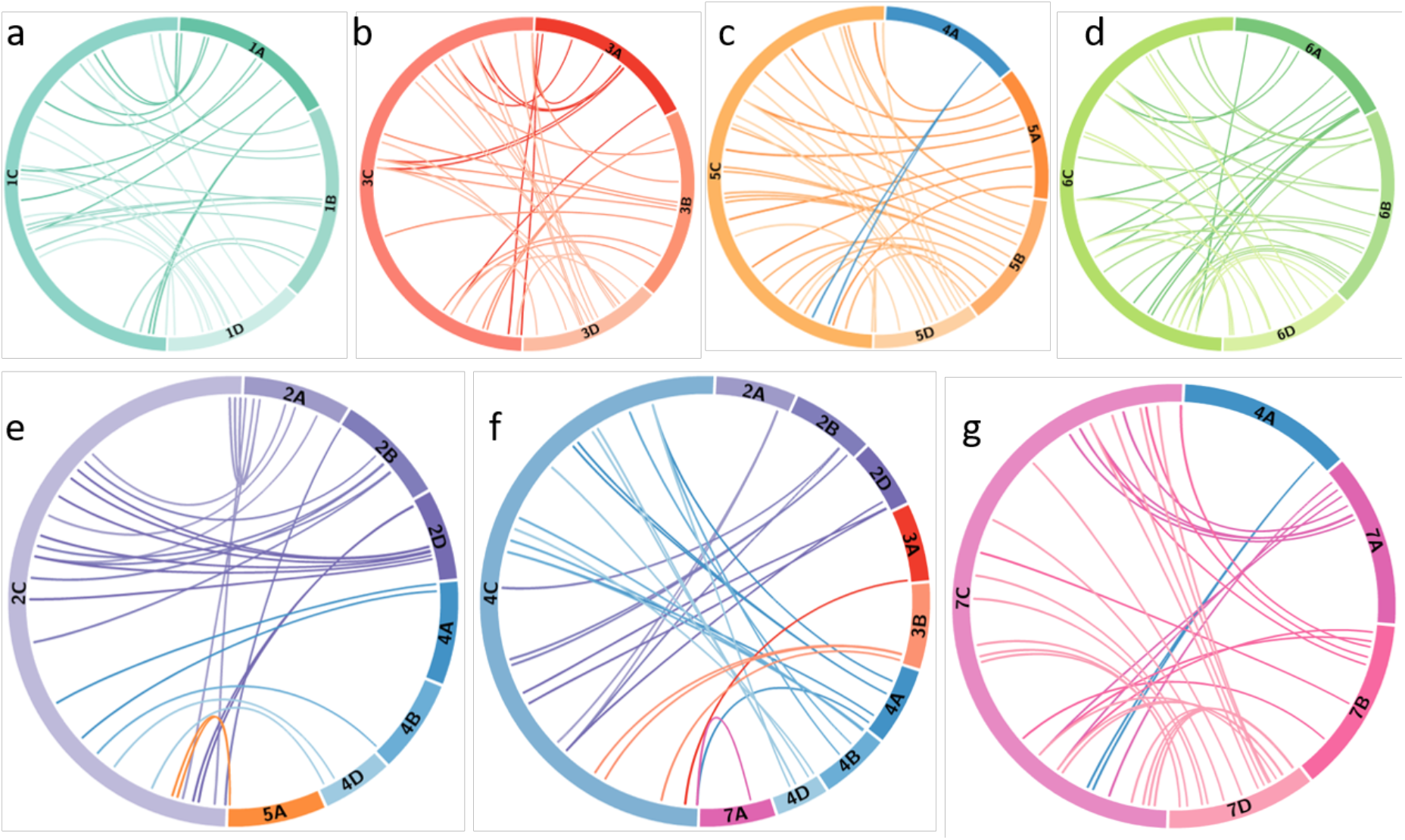
Comparative analysis of *Ae. caudata* chromosome rearrangement compared to wheat chromosomes revealed by physically mapped KASP markers. Circos diagrams of *Ae. caudata* chromosome **a)** 1C, **b)** 3C, **c)** 5C, **d)** 6C, **e)** 2C, **f)** 4C and **g)** 7C. Corresponding markers are linked between their tentative physical order on the *Ae. caudata* C-genome chromosomes and their physical location (Mb) on the orthologous regions on the wheat genome.

KASP marker analysis of the recombinant lines showed that markers WRC0002-0077 on chromosome 1C (Figure 3) corresponding to wheat homoeologous group 1 markers, spanning the short arm and approximately half of the proximal region of the long arm (Figure 4a), detected the *Ae. caudata* segment present in the recombinant chromosome T1AL-1CL.1CS. The spread of these markers across homoeologous group 1 also matched the size of the segment subsequently shown by mcGISH (Figure 2a). KASP markers WRC0095-WRC0142 on chromosome 1C (Figure 3), corresponding to the distal end of the long arm of homoeologous group 1 (Figure 4a), detected the *Ae. caudata* introgression present in T1DS.1DL-1CL subsequently validated by mcGISH (Figure 2b). This suggested that the marker sequences on chromosome 1C were potentially collinear with their counterparts on wheat chromosomes 1A, 1B and 1D.

Chromosome 4C of *Ae. caudata* had KASP markers from wheat homoeologous groups 2/3/4/7 assigned to it (Table 1). KASP markers WRC0387-0875 on chromosome 4C (Figure 3) detected the presence of an *Ae. caudata* segment in recombinant chromosome T3DS.3DL-4CL. However, Figure 4f shows that these markers were physically located on distal end of the long arm of wheat homoeologous group 3 (WRC0387, WRC0390 and WRC0335) with one marker near the centromere on homoeologous group 7 (WRC0875), indicating genomic rearrangements between homoeologous groups 3/4/7 in *Ae. caudata*. Due to the recombinant chromosome being present as a single copy in the introgression line, the KASP markers were not indicative of the wheat chromosome that had recombined with the *Ae. caudata* segment (Figure 2c). McGISH indicated it to be a D-genome chromosome which was subsequently confirmed to be chromosome 3D by FISH analysis (Figure 5a-b). The FISH analysis also showed that one copy of chromosome 1D was missing and hence, the total chromosome number for this line was 41 as mentioned in Table 2.

**Figure 5.**
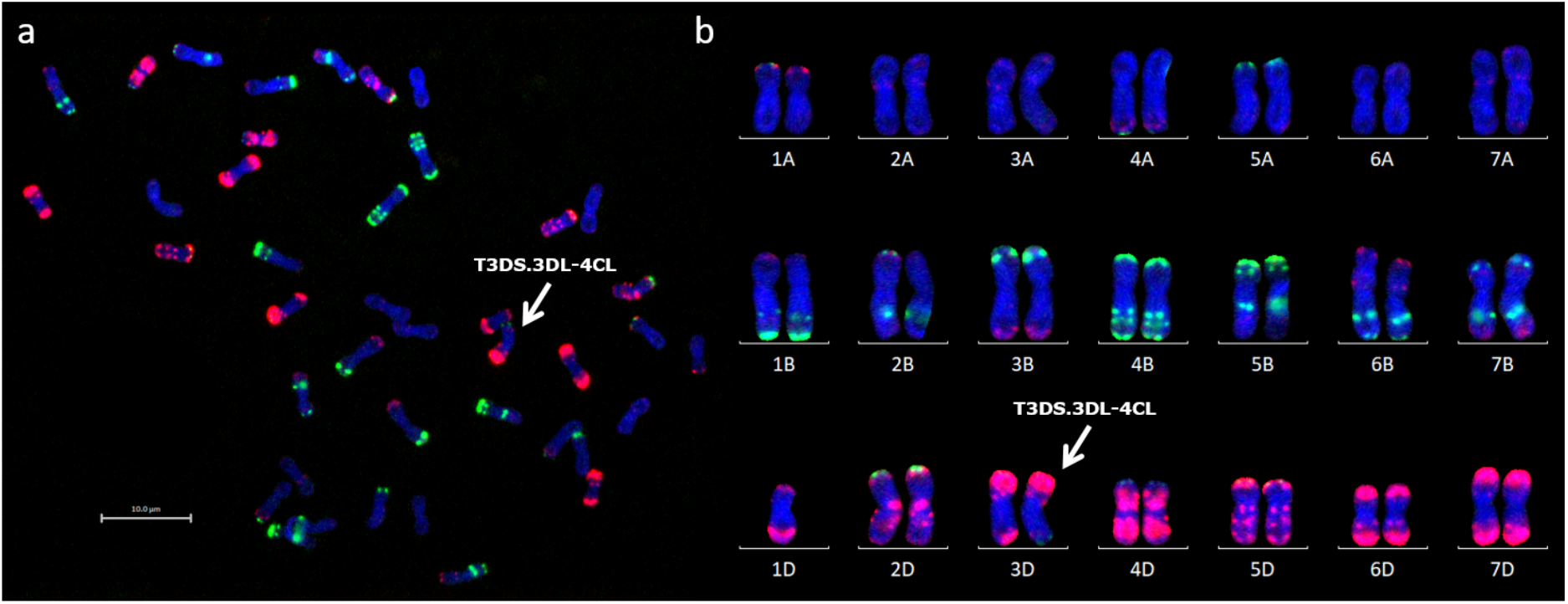
FISH analysis of wheat-*Ae. caudata* introgression line carrying recombinant chromosome T3DS. 3DL-4CL. **a)** FISH image of a root-tip metaphase spread of chromosomes from line BC_4_F_1_-164-2. Arrow indicates recombinant chromosome T3DS. 3DL-4CL. **b)** FISH karyogram of the same spread from line BC_4_F_1_-164-2 showing 41 chromosomes including one copy of recombinant chromosome T3DS. 3DL-4CL and one copy of chromosome 1D. Sites of hybridisation are shown with fluorescence-labelled probes, pSc119.2 (green) and pAs.1 (red). Wheat chromosomes were assigned to the karyogram according to the FISH karyotype established by Tang et al. (2014).

Four recombinants were obtained from chromosome 5C of *Ae. caudata*. KASP marker analysis showed that 21 markers, from WRC0668 to WRC0558 on chromosome 5C (Figure 3) detected all four recombinants. These markers correspond to wheat homoeologous group 5 spanning the small arm and approximately half of the proximal region of the long arm (Figure 4c). However, the four recombinants differed in their size and the wheat genome they had recombined with. For example, T5AL-5CL.5CS#1 and T5DL-5CL.5CS had the same 21 markers detecting the *Ae. caudata* segment but the former had recombined with chromosome 5A of wheat and the latter with chromosome 5D of wheat (Figure 2e and 2f, respectively). Recombinant chromosome T5BL-5CL.5CS had another 3 distal markers (WRC0703, WRC0595 and WRC0662; Figure 3), in addition to the 21, that detected the *Ae. caudata* segment in it and mcGISH analysis showed that it had recombined with the B-genome of wheat (Figure 2g). T5AL-5CL.5CS#2 had the longest segment from chromosome 5C of *Ae. caudata* and had another distal marker (WRC0645; Figure 3), making it a total of 25 markers, that detected this introgression. These markers indicated that recombination happened with chromosome 5A of wheat in this line and was also validated by mcGISH (Figure 2h). These segments assisted in the ordering of markers on chromosome 5C and comparative analysis showed that they were potentially collinear to corresponding sequences on wheat homoeologous group 5 (Figure 4c).

One large *Ae. caudata* segment was detected by 18 KASP markers, from WRC0908 to WRC0937 present on chromosome 7C (Figure 3). These markers indicated the segment to be to be homozygous and recombined with wheat chromosome 7D in the introgression line (Figure 2d). McGISH analysis of the recombinant chromosome T7DL-7CL.7CS showed that the introgressed *Ae. caudata* segment included the short arm and a large part of the proximal region of the long arm of chromosome 7C, with the distal end of the recombinant chromosome being replaced with the distal end of chromosome 7DL (Figure 2d). However, the markers found in this segment were found to be present on the proximal region of the short arm and the whole of the long arm of wheat homoeologous group 7 (Figure 4g). The markers corresponding to the short arm of wheat homoeologous group 7 were not detecting this recombinant chromosome and were thus, physically ordered onto the distal end of chromosome 7C (Figure 3) indicating an inversion in *Ae. caudata* group 7 compared to wheat.

Where markers could not be ordered due to absence of any recombination with wheat, such as for homoeologous group 2, 3, 4 and 6, the markers on chromosome 2C, 3C, 4C (except the distal end of the long arm) and 6C were ordered based on a previous study by Danilova et al. (2017).

## DISCUSSION

*Ae. caudata* is known to carry many useful genes that can be used for wheat breeding. Its utilisation for wheat improvement relies on the development of suitable and reliable molecular markers that can be used for the identification of *Ae. caudata* segments in introgression lines. Friebe et al. (1992) established cytogenetic markers (the C-banding pattern) of *Ae. caudata* chromosomes, Peil et al., (1997, 1998) developed RAPD and SSR markers and Kong et al. (1999) converted cloned specific repetitive DNA from the *Ae. caudata* genome into a marker, all for the detection of *Ae. caudata* in a wheat background. More recently, Gong et al. (2017) developed EST-based markers and Niu et al. (2018) developed SSR markers for the detection of *Ae. caudata* chromatin in a wheat background. In this research, a total of 182 KASP markers, developed in a previous study (Grewal et al., 2020), were assigned to specific *Ae. caudata* chromosomes (Table 1), providing a new rapid detection approach for the quick selection and identification of wheat-*Ae. caudata* introgression lines.

A number of studies have attempted assignment of various types of molecular markers to specific *Ae. caudata* chromosomes by testing them on the Alcedo-*Ae. caudata* disomic addition lines (Schubert and Blüthner, 1992; Schubert and Blüthner, 1995). The variation observed in the results indicates the presence of genomic rearrangements. Recently, studies by Danilova et al. (2017), Gong et al. (2017) and Niu et al. (2018) on the Alcedo-*Ae. caudata* additions lines found that the C-genome chromosomes carried several inversions and translocations but did not agree on the rearrangements carried by each chromosome. In this study, we used the same set of addition lines for the assignment of the KASP markers to the C-genome chromosomes. Genotyping of these addition lines with the KASP markers showed that chromosome B (2C) had markers from wheat homoeologous groups 2/4 assigned to it (Table 1) which agreed with the work done by Danilova et al. (2017). Chromosome G (4C) had KASP markers assigned from homoeologous groups 4/2/3/7. This partially matched the results by Danilova et al. (2017) and Niu et al. (2018) who found that chromosome G had markers from homoeologous groups 4/2/3. Our studies found one marker from homoeologous group 7 assigned to chromosome 4C of *Ae. caudata*. An addition line for chromosome A is absent from this set of addition lines. We assigned KASP markers that were not assigned to chromosomes B-G to chromosome A and found that they were all homoeologous to the group 1 chromosomes of wheat (Figure 4a) which is consistent with previous reports by Danilova et al. (2014, 2017) that determined that chromosome A of *Ae. caudata* is homoeologous to wheat group 1 chromosomes by using FISH with cDNA probes.

To exploit the potential of the C-genome, *Ae. caudata*-derived wheat introgression lines were developed and characterised, primarily, for disease resistance genes. Dyck et al. (1990) found stem rust resistance in crosses between *Ae. caudata* and wheat monosomic 5B lines. Another study mapped *Ae. caudata*-derived leaf rust resistance on chromosome 2AS of wheat (Iqbal et al., 2007) while two *Ae. caudata*-derived powdery mildew resistant quantitative trait loci (QTL) were mapped onto chromosomes 1AS and 7AL of wheat (Weidner et al., 2012). Riar et al. (2012) and Toor et al., (2016) mapped leaf rust and stripe rust resistance, respectively, onto chromosome 5DS of wheat in one wheat-*Ae. caudata* introgression line. Here, we report the generation of 9 whea*t-Ae. caudata* introgression lines (Table 2), of which 6 had stable homozygous recombinant wheat-*Ae. caudata* chromosomes (Figure 1). These lines will be made available for further trait characterisation studies.

Previous work investigating wheat-*Ae. caudata* introgression lines did not attribute the disease resistance QTLs they found in these lines to a specific *Ae. caudata* chromosome. Gong et al., (2017) and Niu et al. (2018) evaluated the Alcedo-*Ae. caudata* addition lines for agronomic traits such as resistance to rust and powdery mildew diseases. Addition lines D (6C), F (3C), and G (4C) showed some resistance to powdery mildew isolates (Niu et al., 2018) while addition line E (7C) showed a higher resistance to powdery mildew (Gong et al., 2017; Niu et al., 2018). Gong et al. (2017) suggested chromosome D (6C) carried leaf rust resistance while Niu et al. (2018), suggested that chromosome B (2C) exhibited leaf rust resistance at a level comparable to the *Ae. caudata* parent. While Gong et al. (2017) suggested that chromosome E (7C) might carry some stem rust resistance, Niu et al. (2018) found no stem rust resistance in the addition lines. However, addition lines C (5C) and D (6C) had been previously identified as resistant to the Ug99 race group of the stem rust pathogen (*Puccinia graminis* f. sp. *tritici*; Xu et al., 2009). Consequently, a new gene for Ug99 resistance was introgressed from chromosome D (6C) into wheat (Xu et al., 2017). Investigations into grain quality traits showed that chromosome E (7C) or F (3C) could increase protein and wet gluten content when introduced into wheat (Gong et al., 2017). In this study, we have produced four wheat-*Ae. caudata* recombinant lines with wheat from chromosome C (5C) and one from chromosome E (7C) (Table 2), among others, which could potentially carry disease resistance and grain quality QTLs.

Genotyping of the back-crossed population with the KASP markers showed that the markers were not only able to detect *Ae. caudata* in the wheat background but also indicated whether a line had a homozygous segment and which wheat chromosome had recombined with *Ae. caudata* in a homozygous line. Eight out of nine introgression lines had a single type of recombinant chromosome (Table 2). The ninth line was homozygous for two of these recombinant chromosomes. As shown in Figure 2 (bottom), the KASP markers were able to detect whether a line was homozygous (red and green marker segments) or heterozygous (all red marker segments). A wheat chromosome that had green marker segments, representing homozygous *Ae. caudata* alleles, was shown to be involved in the recombinant chromosome. The genotyping results were also validated by mcGISH analysis of the recombinant lines (Figure 2; top).

Genotyping of these recombinant lines helped to physically order some of the KASP markers along the *Ae. caudata* chromosomes (Figure 3). For example, the recombinant line with chromosome 7C showed that markers homoeologous to the distal end of the short arm of wheat homoeologous group 7 were potentially present on the distal end of the long-arm of chromosome 7C in *Ae. caudata* (Figure 4g). This agrees with a previous report by Danilova et al. (2017). Taking this into account and the other marker assignments discussed above, we concluded that our results fit well with the conclusions by Danilova et al. (2017). Thus, for the regions of the C-genome where no recombination was obtained, the markers were ordered according to Danilova et al. (2017; Figure 3) and compared to the wheat genome (Figure 4) which showed all the potential inversions and translocations found in the C-genome of *Ae. caudata*.

Chromosomes 1C, 5C, 6C and 7C had maintained macro-synteny with wheat while chromosomes 1C and 5C were collinear with wheat and chromosomes 6C and 7C had inversions (Figure 4). In contrast, chromosomes 2C, 3C, and 4C had inter-chromosomal translocations and inversions (Figure 4). Due to the 4/5 translocation in wheat’s A-genome, Figures 4 showed markers corresponding to wheat chromosome 4A for *Ae. caudata* chromosome 5C (Figure 4c) and markers corresponding to wheat chromosome 5A for *Ae. caudata* chromosome 2C (that had a 2/4 translocation; Figure 4e). The macro-synteny and macro-collinearity with wheat could potentially be the reason we were able to obtain 6 recombinant chromosomes from *Ae. caudata* chromosomes 1C and 5C (Table 2; Figure 2). Even though the majority of our results agreed with those of Danilova et al. (2017), we did not find any markers from wheat homoeologous group 7 translocated onto chromosome 6C (Table 1 and Figure 4f). This may be due to differences in the *Ae. caudata* accessions used in the two studies.

Our work also found one marker from wheat homoeologous group 7 assigned to chromosome 4C [not found by Danilova et al. (2017)], present in recombinant chromosome T3DS.3DL.4CL along with 3 markers from wheat homoeologous 3. Where a line was heterozygous, the KASP markers were not able to indicate the wheat chromosome involved in the recombinant chromosome. McGISH analysis of this line had shown that the introgression line was heterozygous for the segment and a wheat chromosome from the D-genome was involved. Figure 4f indicated that markers detecting the segment belonged to wheat homoeologous groups 3/7. To confirm whether the recombination in this chromosome took place with wheat chromosome 4D (since the *Ae. caudata* segment was from chromosome 4C) or wheat chromosome 3D (since the majority of the KASP markers were placed on the distal end of wheat homoeologous group 3), the line was subjected to FISH analysis (Figure 5). FISH karyotyping of this line indicated that the chromosome involved in the recombination event was wheat chromosome 3D (Figure 5b). This confirmed the translocation of wheat homoeologous group 3 onto chromosome 4C, possibly via a meiotic crossover between homologous regions of the translocation (Schubert, 2007).

In contrast to most diploid and allopolyploid Triticeae, that have preserved chromosome macrostructure, rye (*Secale cereale*), *Aegilops umbellulata* and *Ae. caudata* genomes are highly rearranged (Danilova et al., 2017; Devos et al., 1993). When collinearity between the homoeologous wheat and wild relative chromosomes is interrupted, as a consequence of evolutionary chromosome rearrangements in the wild relative genomes (Devos et al., 1993; Zhang et al., 1998), meiotic chromosome pairing is reduced or absent. However, the recombinant chromosome, T3DS.3DL.4CL, along with a recombinant from chromosome 7C, T7DL-7CL.7CS (which has an inversion from the short arm onto the long arm compared to wheat) developed in this work show that it is possible to get wheat-wild relative recombination from wild relative chromosomes that are highly rearranged with respect to wheat.

Genomic rearrangements can be caused in Triticeae interspecific crosses due to gametocidal (Gc) genes, which cause chromosome breakage in the gametes that lack them (Maan, 1975; Tsujimoto, 1995). Gc genes were found in chromosome 3C of *Ae. caudata*-derived allotetraploids and other Aegilops species (reviewed by Endo, 2007). Some Gc genes can induce mild chromosome changes, hence structurally rearranged chromosomes may be retained (Endo, 2007; Friebe et al., 2000). Apart from the lack of the *Ph1* gene in the F_1_ interspecific hybrids, this Gc action could potentially also be responsible for the translocations between wheat and *Ae. caudata* chromosomes shown in this work. It is said that the intensity of the Gc action is variable, from lethal to semi-lethal and is affected by the genotypes of the common wheat lines where the Gc chromosomes are present (Endo, 1990). Some Gc chromosomes, such as 4S^sh^ from *Aegilops sharonensis*, can cause complete sterility in gametes without themselves in ‘Chinese Spring’ wheat and therefore are exclusively transmitted to the offspring. (Grewal et al., 2017; Miller et al., 1982). Chromosome 3C exerts severe Gc action in Chinese Spring but has mild, semi-lethal action in some other cultivars, implying the presence of an inhibitor gene in those cultivars. and suggesting that it would not be preferentially transmitted in these other wheat cultivars (Endo and Gill, 1996). Our work does not show any preferential transmission of chromosome 3C in the back-crossed population involving wheat cultivar Paragon (data not shown), which is consistent with this theory. It is also possible that the *Ae. caudata* accession used in this study does not possess any Gc action and all recombinant chromosomes were obtained due to homoeologous recombination during meiosis. Six out of the eight recombinant chromosomes obtained in this work were between homoeologous linkage groups, i.e. chromosomes 1C of *Ae. caudata* with chromosomes 1A and 1D of wheat (Table 2) whereas any Gc action would have potentially resulted in random translocations.

Wheat-wild relative introgressions play an important role in wheat improvement. Production of novel wheat-wild relative recombinant chromosomes, particularly for disease resistance, has always been the research hot topic. However, genes on wild relative chromosome segments that do not compensate for the loss of wheat genes have a negative effect on agricultural performance of the wheat–wild relative introgression lines (Friebe et al., 1996; Sears and Gustafson, 1993). Therefore, better knowledge of the genome organisation of the wild relative species is essential before embarking on the time-consuming task of producing compensating translocations for commercial agriculture. In this work, we produced KASP markers that proved to be valuable tools to detect *Ae. caudata* chromatin in a wheat background. In addition, these KASP markers have shed further light on the structural rearrangements present in *Ae. caudata*. We have also produced nine wheat-*Ae. caudata* recombinant lines of which six have stable homozygous introgressions that can be used for further trait analysis studies.

## AUTHOR CONTRIBUTIONS

SG, MO, JW, CY, SHE, DS, SA, IPK and JK carried out the crossing programme. PI set up the high-throughput genotyping platform and developed the protocol. MO and JW performed the *in situ* hybridisation experiments and the genotyping of wheat-*Ae. caudata* lines. SG analysed the genotyping data, assigned the markers to the *Ae. caudata* chromosomes and performed the comparative studies. SG, MO, IPK and JK conceived and designed the experiments. SG wrote the manuscript with assistance from MO, JK and IPK. All authors read and approved the final manuscript.

## ACKNOWLEDGEMENTS

This work was supported by the Biotechnology and Biological Sciences Research Council (Grant No. BB/P016855/1) as part of the Developing Future Wheat (DFW) programme. The funding body played no role in the design of the study and collection, analysis, and interpretation of data and in writing the manuscript.

